# Why do bats fly into cave doors? Inattentional blindness in echolocating animals

**DOI:** 10.1101/2024.10.16.618711

**Authors:** Nikita M Finger, Keegan E Eveland, Xiaoyan Yin, Cynthia F Moss

## Abstract

Echolocating bats can navigate complex 3D environments by integrating prior knowledge of spatial layouts and real-time sensory cues. This study demonstrates that inattentional blindness to sensory information undermines successful navigation in Egyptian fruit bats, *Rousettus aegyptiacus*, a species that has access to vision and echolocation to traverse natural environments. Bats flew over repeated trials to a perch at a fixed location in the light, allowing them to navigate using both vision and echolocation. The experiment was then repeated in the dark to exclude the bat’s use of vision. The perch was subsequently displaced by either 15 or 30 cm in one of six different directions (up, down, left, right, front, back). Echolocation behavior was recorded using a 25-channel microphone array, while flight paths were tracked using 13 motion capture cameras. The directional aim of echolocation clicks served as a metric for the bat’s spatial attention to locations in their environment. In the light, bats modified their flight paths to successfully land on a perch that was moved 15 cm but surprisingly, often failed to land on it when displaced by 30 cm. In the dark, bats often failed to land on the perch after it was moved by only 15 cm. Landing failures suggest that learned spatial priors invoked inattentional blindness to changes in the environment, which interfered with successful navigation. In both the light and dark, when bats failed to land on the perch at its new location, they directed their attention toward the original perch position. Performance differences in the light and dark suggest that the bat’s attentional spotlight may be narrower when it relies on echolocation than vision. To our knowledge, these findings provide the first evidence of inattentional blindness in a flying echolocating animal, demonstrating that spatial priors can dominate sensory processing during navigation.

## Introduction

Spatial navigation builds on the integration of prior knowledge with real-time sensory data to guide movement and predict spatial dynamics (1–4) Spatial priors—expectations or assumptions about the environment formed through repeated experiences—play a crucial role in this process. Spatial priors are probabilistic internal models that guide perception and action by directing attention to events or spatial locations based on instructions or past encounters (5–7). Sensory cues from vision, hearing, touch, and smell provide additional information to guide navigation, particularly in novel environments (8). However, the relative weighting of spatial priors and sensory information depends on an organism’s behavioral state and the specific demands of a task (9).

In stable and familiar environments, reliance on spatial priors reduces cognitive load associated with processing sensory information, enabling more efficient navigation (3,10,11). Research by Zhang et al. (2022) demonstrated that attentional biases are optimized according to learned spatial probabilities in a given environment. Their study showed that when a target appeared more frequently in certain locations, participants’ attention became biased towards those locations, leading to faster and more accurate responses. This reliance can be a limitation, as a subject continues to depend on priors at the expense of detecting changes in the environment leading to navigation errors. This phenomenon, known as inattentional blindness (IB), occurs when an individual fails to notice new or unexpected stimuli, because their attention is focused on other aspects of the environment. Consequently, stimuli in unexpected locations are often overlooked (12,13).

Inattentional blindness has been attributed to the adjustment of a spatial priority map derived from statistical learning, highlighting that learned spatial priors can both facilitate efficient distribution of attention and lead to perceptual oversights (5,7,14). In such cases, even salient objects or stimuli can be overlooked, not due to sensory processing limitations, but because attention is anchored to expected features (13). Whether inattentional blindness reflects perceptual or memory failures remains a topic of debate (14–16).

Bats are compelling model organisms to investigate the interplay between spatial priors and sensory processing in navigation tasks. Many bat species rely on echolocation, a sensory modality that operates effectively in darkness for small-scale navigation (17). Echolocation relies on comparisons made between call production and echo reception (17–19). Despite their sophisticated echolocation abilities, bats do not always rely on echo returns to guide their navigation, as highlighted by historical observations (18,20–24). In the 19th century, a curious phenomenon was noted following the installation of safety doors in caves, which led to an increase in bat fatalities from collisions with these doors. The most notable case occurred in Wyandotte Cave, home to a large population of little brown bats, *Myotis lucifugus*, that slammed into newly installed cave doors, resulting in thousands of bat deaths (21). Other studies investigated this phenomenon and confirmed that bats were indeed producing echolocation calls during navigation failures in familiar environments (18,23), but it is not known whether all bats in large groups produce echolocation calls or where they direct their sonar when they enter caves. This leaves unanswered questions about the bat’s spatial attention during navigation failures (17,18,23,25).

The Egyptian fruit bat, *Rousettus aegyptiacus*, is a particularly well-suited species to study the relative weighting of spatial priors and sensory cues in navigation. It is documented that Egyptian fruit bats rely on cognitive maps to travel to and from feeding sites (26–29). This bat species uses both scotopic vision (30,31) and lingual echolocation (18,30,32) for orientation, setting them apart from many other echolocating bat species (33). Unlike most echolocators that produce sonar vocalizations, Egyptian fruit bats produce echolocation clicks with the tongue (32–36). These clicks are produced in pairs and directed off-axis from the midline, allowing the bat to use the maximum slope (“edge”) of the beam for target localization (34,35). Importantly, the directional aim of the bat’s sonar clicks can be used as a metric of the bat’s spatial attention to objects in its environment.

The present study revisits the decades-old question of why bats collide with obstacles (18,21) and provides compelling evidence for inattentional blindness in flying mammals. Further, these findings contribute to the ongoing debate over whether inattentional blindness reflects perceptual limitations or memory failures. By combining a microphone array with high-speed stereo video recordings of Egyptian fruit bats navigating in both light and dark, this study quantified where bats directed their echolocation clicks, and thus their sonar attention in the environment. We hypothesized that bats operating in familiar and unchanging environments establish spatial priors, which can override sensory cues and give rise to inattentional blindness. We predicted that bats trained to fly to a landing perch in a fixed location will fail to find the perch after it has been moved, as their attention is directed at the location governed by learned spatial priors.

## Results

Eight bats were trained in this study (four trained in the light, four trained in the dark). Four bats trained in the light reached successful landing criteria of 80% (two males, two females). One female bat died of natural causes before a complete data set could be collected. Four bats trained in the dark failed to reach the 80% success landing criteria after a combined total of 416 trials. Bats initially trained in the light quickly reached testing criteria again in the dark (S1 Table). In total, 387 trials were conducted in the light and dark, across three bats, including 50 probe trials where the perch was moved (S2 Table).

### Flight Trajectories

Bats’ flight trajectories included direct successes (DS), back-and-forth failures (BFF), back-and-forth successes (BFS), and direct failures (DF) (Fig 1). Example flight trajectories are shown for different perch displacements in the light in Fig 2. A multinomial logistic regression revealed significant effects of lighting and perch location on flight trajectories with several key contrasts showing differences in flight trajectories (GLMM; χ35,343 = 175). The model converged with a final deviance of 316.82 and an AIC of 693.64. Bats showed similar flight patterns to the standard perch location and to the perch moved 15 cm in the light (p>0.05 for DS, BFF, BFS, and DF). When the perch was displaced by 30 cm in the light, bats made fewer direct successful landings compared to both the standard (p<0.001) and 15 cm perch-moved conditions (p=0.048). Bats failed to land on the perch most often when it was moved by 30 cm in the light compared to when it was moved by 15 cm in the light and dark (p<0.05). In standard trials, bats in the dark made more repeated approaches to the perch before landing than in the light (p<0.01). In contrast to the standard location in the dark, bats showed no direct landings after the perch was moved by 15 cm (p<0.001). Bat performance in the light and dark 15 cm perch-moved conditions was also significantly different (p<0.001) (Fig 2). There were no significant differences in flight patterns among individual bats (p>0.05 for all bats), although Bat 2 exhibited a tendency to exhibit more back-and-forth successful landings compared to direct success across all conditions (S3 Fig).

**Fig 1:**
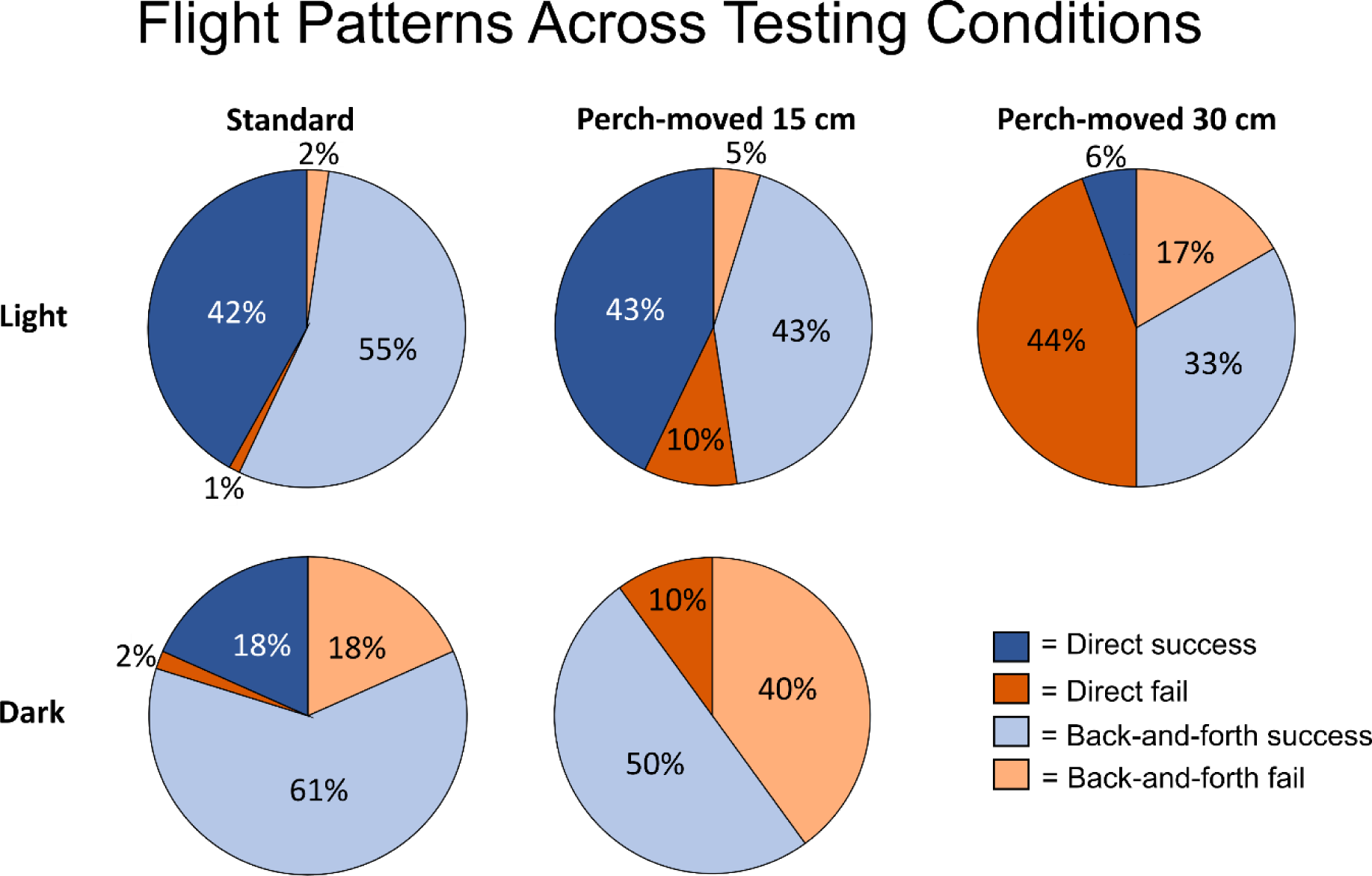
Flight patterns across different perch displacements, and lighting. Egyptian fruit bats’ flight approach and landing outcomes under different perch displacements (standard, perch-moved 15 cm, and perch-moved 30 cm) and lighting conditions (light and dark). Approach and landing performance are categorized into four distinct outcomes: direct success, direct failure, back-and-forth success, and back-and-forth failure.

**Fig 2:**
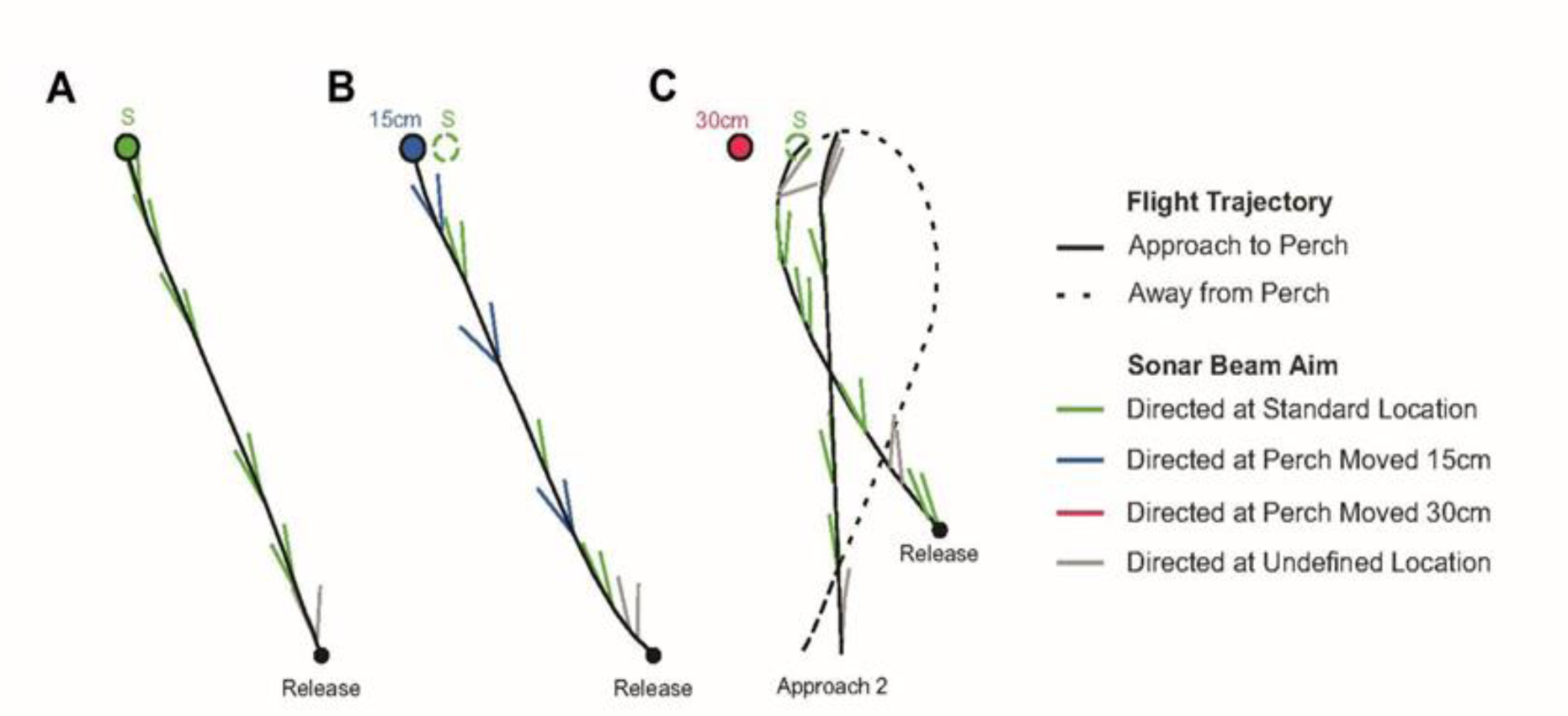
Schematic of example flight trajectories and sonar beam aim. Example flight trajectories and directional aim of sonar clicks when the perch was positioned in three different locations in the light: A standard location, B perch-moved 15 cm, and C perch-moved 30 cm. Approaches to the perch are illustrated by solid black lines, while departures are shown by dashed black lines. Echolocation clicks are represented by colored lines along the flight trajectory. Different color lines indicate sonar beam direction derived from analysis of each click-pair. Lines indicate where the click-pairs are directed: green indicates the standard perch location, blue indicates the perch location moved 15 cm, and gray indicates an undefined location outside of the perch locations. Note that echolocation click pairs were sometimes aimed at the standard location when the perch was moved from that location (dashed circles) in the perch-moved 15 cm trial and consistently in the perch-moved 30 cm trial. Notably, in this example, the bat produced no sonar clicks directed at the perch when it was moved by 30 cm from the standard location, hence there are no sonar beam clicks denoted in red in panel C.

### Sonar Click-Rate

When the perch was at the standard position, bats increased their click-rate near the perch as they prepared for landing (37). Sonar click-rate is therefore reported for a) the entire approach to the perch and b) as the bat neared the perch (<1.5 meter from the perch). S5 and S6 figures show sonar click rate across different perch displacement magnitudes (standard vs perch-moved 15 cm vs perch-moved 30 cm), lighting conditions (light vs dark), and landing conditions (landed vs didn’t land). There were no significant differences in click-rates across different perch displacements (GLMM: F₃, ₁₆₀=0.127, N=176, p=0.944; GLMM: F₃,₁₆₀=1.954, N=176, p=0.123) (S5 Fig). Lighting was a significant factor in predicting click-rate (GLMM: F₁, ₁₆₀=16.421, N=176, p<0.01; GLMM: F₁, ₁₆₀=10.583, N=176, p<0.01). Landing was only a significant predictor of click-rate near the perch (GLMM: F₁, ₁₆₀ = 13.814, N=176, p < 0.001) but not over the entire approach to the perch (F₁, ₁₆₀ = 2.024, N=176, p = 0.157). There were no significant interactions between perch displacement, lighting, and landing (GLMM: F₂, ₁₆₀=0.921, N=176, p=0.400, GLMM: F₂, ₁₆₀=0.709, N=176, p=0.494). In standard trials, bats produced higher click-rates in the dark than in the light over the entire distance to the perch (Friedman ANOVA: χ²(2) =114.920, N=88, N=3, p<0.001): Bat 01 (Wilcoxon Signed Rank Test: z=2.203, N=13, P=0.028), Bat 02 (z=2.982, N=12, p<0.01), and Bat 03 (z=3.622, N=17, p<0.01).

### Landing Success and Beam Directing Behavior

The bats’ landing success was significantly influenced by both lighting conditions and perch movement. When the perch was moved, the bats’ sonar beam directing behavior revealed that they continued to direct their spatial attention to the standard perch location (where the perch was absent), which corresponded with failures to find the perch at its new position. Generalized Linear Mixed Models (GLMM) analysis showed that lighting condition (p < 0.01) and perch displacement (p < 0.01) were significant predictors of landing success, with the model explaining 88.9% of the variance (GLMM: F₃₆,₃₂₈ = 0.736). The analysis showed no significant relationship between landing success and perch displacement direction (left, right, up, down, front, back) (p = 0.324), landing success over time (p = 0.920), release location (left, middle, right) (p = 0.888), individual bat (p = 0.093), or interactions between these variables (Unequal-N p > 0.05 for all interactions). Therefore, data for different perch movement directions and release locations were combined, as these factors did not significantly affect landing success.

In the light, bats landed successfully on the perch in 86% of the trials after it was moved 15 cm, a landing success rate that was not significantly different from the standard perch location (p=0.810). In contrast, bats landed successfully in only 39% of the trials when the perch was moved by 30 cm in the light. This decrease in landing success after the perch was moved 30 cm was statistically significant compared to both standard (p<0.01) and perch-moved 15 cm conditions (p<0.01) (Fig 3). In the dark, bats showed a decrease in landing success to 50% when the perch was moved by 15 cm, compared to 80% when it was in the standard location (GLMM: F36,328 = 0.736; Unequal-N p<0.01). The bat’s landing performance was significantly lower when the perch was moved by 15 cm in the dark (30% reduction from standard location) (p=0.012) (Fig 3).

**Fig 3:**
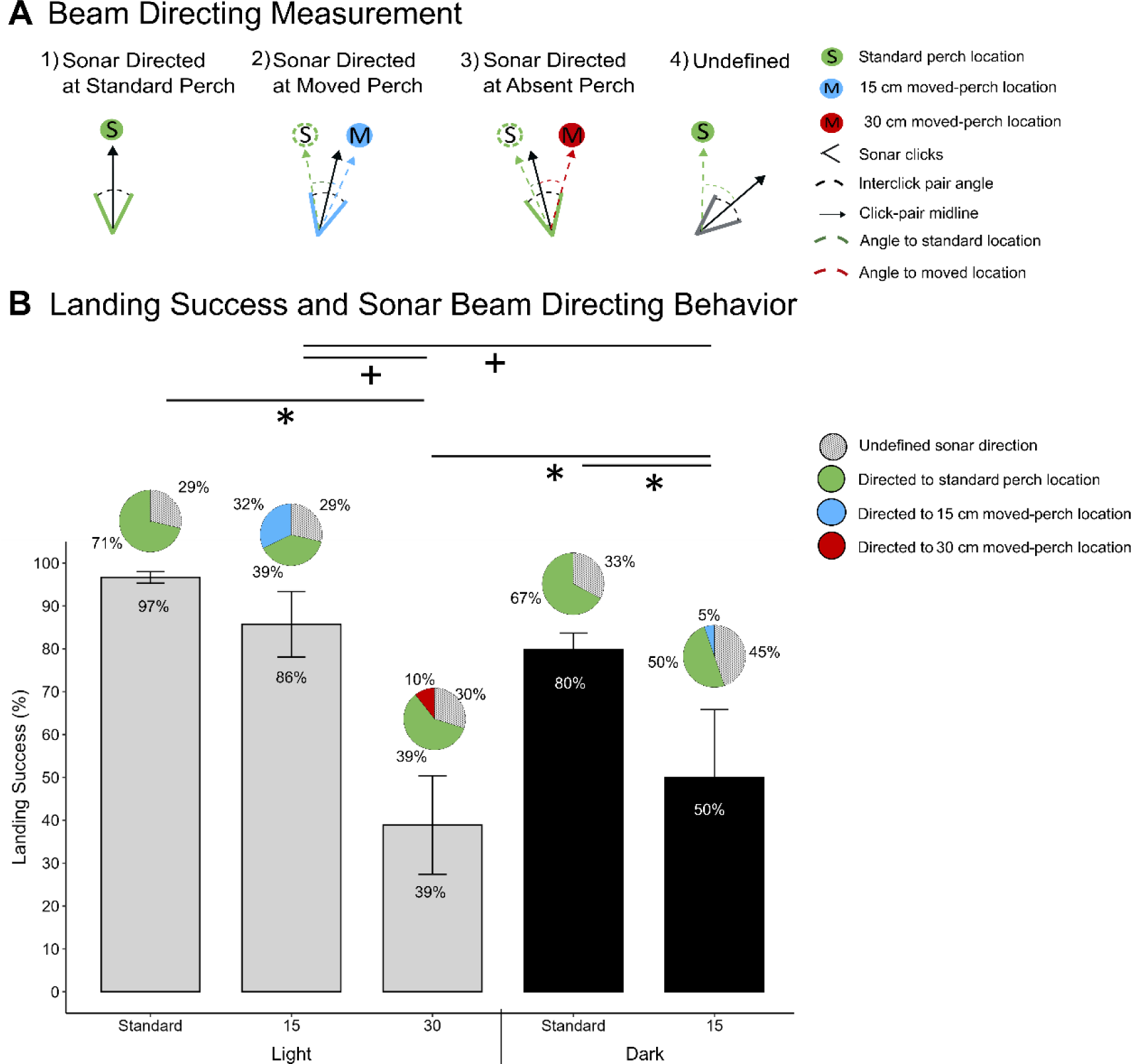
A: Metric of sonar-guided attention: The directional aim of echolocation clicks. Schematic of sonar click-pairs with angular offsets relative to the perch in different conditions. The midline of a click-pair (black arrow) in 1) is directed towards the standard perch location (S, shown in green). The midline of a click-pair in 2) is directed towards the 15 cm moved perch location (M, shown in blue). The midline of the click-pair in 3) is directed towards the standard perch location (S) in a 30 cm perch-moved trial. The midline of a click-pair directed outside the two locations of interest, illustrated in panel 4), were categorized as undefined, based on a ±10 deg tolerance to the nearest perch location (shown in gray). **B. Landing behavior and sonar beam aim reveals inattentional blindness in perch-moved trials.** Bar graph of the landing success (± SEM) of bats when the perch was either at the standard location, moved 15 cm, and moved 30 cm in the light and dark. Pie charts depicted above performance plots for each condition, show the aim of the sonar click pair towards the standard perch location (green), the perch-moved by 15 cm (blue) or 30 cm (red), or in an undefined direction (striped) within that condition. Note when the bat directed its sonar beam toward the standard perch location after the perch was moved by 15 or 30 cm (shown in green), it directed its sonar to a location where the perch was absent. Significance differences between landing success across conditions is denoted by asterisks; significance between sonar beam directing behavior across conditions is denoted by pluses.

The bats’ sonar beam directing behavior varied with the magnitude of perch displacement (15 cm versus 30 cm) and lighting conditions, revealing differences in their ability to locate the perch under different perch-moved conditions. Larger perch displacements (30 cm) and dark conditions led to fewer clicks directed at the moved perch location with more failures to land. There were significant differences in the number of echolocation click pairs directed at the new perch location (Kruskal-Wallis chi-squared = 6.714, df = 2, p = 0.035) and a non-significant trend in the number of echolocation click pairs directed at undefined locations (Kruskal-Wallis chi-squared:5.48, df=2, p=0.064) in the perch-moved trials. There were no significant differences in the number of click pairs directed at the standard perch location (Kruskal-Wallis chi-squared = 1.8422, df = 2, p = 0.398) in the perch-moved trials. Even when the perch was absent from the standard location, bats continued to direct their sonar attention there, directing between 39% and 60% of their clicks at the standard location in perch-moved trials (Figs 1 and 3, S4 Table).

In the light, when the perch was moved 15 cm, bats directed clicks toward the moved perch location, dividing their sonar attention between the standard and moved perch positions (Fig 3, S4 Table). They directed 32% of clicks toward the moved perch location and 39% toward the standard location (Fig 3, S4 Table). This behavior differed significantly from the perch-moved 30 cm in the light condition, where bats directed only 10% of clicks toward the moved perch location (Dunn’s post-hoc test: z = –2.213, p = 0.0202). It also differed significantly from the perch-moved 15 cm in the dark condition, where bats directed only 5% of clicks toward the moved perch location (Dunn’s post-hoc test: z = –2.410, p = 0.0239; Fig 3, S4 Table). There were no significant differences in the number of clicks directed toward the moved perch location between the perch-moved 30 cm in the light and perch-moved 15 cm in the dark conditions (Dunn’s post-hoc test: z = 0.478, p = 0.3162). In both these conditions bats failed to inspect the perch at the new location, and instead directed their sonar attention to the standard location (where the perch was previously located), resulting in fewer successful landings on the perch (Fig 3, S4 Table).

When the perch was at the standard location or in perch-moved trials, clicks directed at undefined locations represented between 29% and 45% of the total clicks. When the perch was moved by 15 cm in the dark, bats directed a larger percentage (45%) of clicks at undefined locations in the room, compared to 30% in the perch-moved 30 cm in the light condition (Dunn’s Post-hoc test: z=0.011, p=0.0333).

## Discussion

To our knowledge, the findings of this study offer the first evidence for inattentional blindness in echolocating bats and address the longstanding question of why bats exhibit navigation failures in familiar environments. In the light, Egyptian fruit bats often failed to land on a perch that had been displaced by 30 cm from its familiar location, despite having access to both visual and echo information signaling the perch’s new position. In the dark, landing failures occurred following a perch displacement of only 15 cm. In both light and dark conditions, when bats missed the perch, they consistently directed their sonar toward the area where the perch had previously been located, providing a metric of their inattention to sensory cues from the moved perch. Metrics of inattentional blindness are consistent with the hypothesis that spatial priors override new sensory cues, particularly in the dark when the spotlight of attention may narrow.

### Learned Spatial Priors in Bats

Spatial priors refer to knowledge or expectations about the location of objects or events in the environment based on experience or learned information. In the context of attention and perception, spatial priors help an organism anticipate where important stimuli are likely to appear and allocate attention more efficiently (5–7). For example, when regularly navigating a familiar environment, such as one’s home, individuals rely on spatial priors to anticipate where furniture, doors, or obstacles are located. This gives rise to a form of Bayesian inference, where prior knowledge (spatial priors) combined with current sensory data, is used to make predictions about the environment (38). In familiar settings, bats navigate complex three-dimensional environments by relying on learned spatial information, reducing the need for continuous perceptual updates and instead, executing pre-planned movements based on expected encounters in the environment (18,20,23,25). If the environment is stable, spatial priors greatly reduce cognitive load. However, when the environment changes unexpectedly, reliance on spatial priors can result in errors (12,13,39) as the bats exhibited in this study.

The results of the present study provide empirical evidence that bats develop spatial priors when navigating in a stable environment, and these priors can give rise to inattentional blindness. In the light, where bats had access to both vision and echo information, they failed in 61% of trials to find the landing perch after it was moved by 30 cm from its original position and instead flew to the standard perch location, suggesting that spatial priors interfered with detection of the perch’s new position. Of the 39% of trials where bats successfully landed on the perch displaced by 30 cm in the light, only 6% were direct successful landings, with the remaining 33% showing multiple attempts before bats finally reached the perch. Among the failures, 44% were direct failures and 17% were back-and-forth flights preceding failures, where the bats made multiple attempts but ultimately did not land on the perch. In the dark bats missed the perch 50% of the time when it was displaced by only 15 cm, and when they did land, none of these landings were on the first approach to the perch.

Past research has demonstrated that previously learned spatial information drives navigational behaviors and may eclipse sensory cues. For instance, Jensen et al. (2005) found that big brown bats, *Eptesicus fuscus,* learned to navigate through an opening in a mist net, cued by an adjacent landmark, and crashed into the net on catch trials where the landmark was displaced from the net opening (23). Helversen and Winter (2003) observed Glossophagine bats repeatedly returning to the previous location of a nectar feeder, despite its relocation (22). Similarly, historical accounts by Blatcheley (1896) documented little brown bats, *Myotis lucifugus*, flying into newly placed doors in previously open flyways, often with fatal consequences (21). Griffin (1958) reported that when bats were trained to navigate through a gap to reach food they repeatedly crashed into the wall when the gap was moved to a new location (18). Neuweiler and Möhres (1967) observed that greater false vampire bats, *Megaderma lyra*, maintained a pose required to squeeze through a previously narrow passage, even after the passage was widened (20).

Thiele and Winter (2005) discovered that nectar-feeding bats use a hierarchical strategy to relocate food targets, relying on spatial memory over cue-directed search (40). Stich and Winter (2006) showed that bats struggle to adjust their behavior in object discrimination when they were placed in new spatial contexts (41). In their study, the researchers used a two-alternative forced choice setup, where bats had to choose between two food options. These food sources were placed in specific locations, but the availability of food at each location changed unpredictably. Despite the irregularity of food rewards, bats strongly associated specific locations with food and continued to rely on spatial cues rather than adjusting their behavior based on other information, such as the changing patterns of food availability. The authors credited the bats’ difficulty to adjust their behavior to their strong dependence on spatial memory, but they did not measure where the animals directed their sonar, leaving a gap in understanding the underlying cognitive and perceptual processes.

### Sonar-guided Attention Confirms Inattentional Bias in Egyptian Fruit Bats

Measurement of the sonar beam directing behavior of free flying Egyptian fruit bats in the present study revealed important insights into navigation failures. When bats missed the moved perch, in the light and dark, they consistently directed their sonar towards the perch’s original location in the room, despite its absence. Their click-rates remained unchanged across standard and moved perch trials. The lack of significant differences in click-rates between the standard and moved perch positions suggests that the bats did not detect the perch in its new location, because their attention was directed towards the standard location, or a learned spatial prior. This was confirmed by measurements of the bat’s sonar beam directing behavior. This finding demonstrates that bats failed to detect unexpected changes in the perch location, an illustration of inattentional blindness in bats.

The phenomenon of inattentional blindness is well documented in humans. For example, there are many reports of walkers, drivers and pilots who fail to notice objects within their visual field of view when engaged in other tasks, such as cell phone conversations (42–44). One of the most striking demonstrations of this effect is the “gorilla experiment” by Simons and Chabris (13). In this experiment, participants were asked to watch a video of people playing basketball and count the number of passes made by players in white shirts. Surprisingly, nearly half of the participants failed to notice a person in a gorilla suit walking directly through the scene. This occurred because the subject’s attention was focused on the task of counting passes, and they completely overlooked the unexpected stimulus. Similarly, in our study, the bats were blind to the new perch location, because they directed their attention to an adjacent region where the perch was typically located.

### Inattentional Blindness through the Lens of an Attentional Spotlight

Early models of attention offer explanations of the processes mediating inattentional blindness. Broadbent’s filter model, introduced in Perception and Communication (45), expanded on William James’s early concept of selective attention introduced in *The Principles of Psychology* (46). Broadbent proposed that our cognitive systems have a limited capacity to process information. His model suggests that when multiple stimuli are present, the mind employs an early selective filter, allowing only one source of information—based on physical characteristics like location or pitch—to pass through for further processing. Posner’s Spotlight Model later expanded on this notion, likening attention to a flashlight. According to this model, attention has a highly focused center where perceptual details are clear, a fringe where information is perceived less distinctly, and an outer margin where stimuli are largely unnoticed (39).

In familiar environments, spatial priors guide the spotlight of attention toward areas where obstacles or targets, such as a landing perch, are expected. This selective focus enables efficient navigation by filtering out irrelevant stimuli in the fringe and margin, thereby reducing cognitive load and streamlining movements (39). However, this attentional focus can interfere with the detection of objects outside the focal region, producing inattentional blindness. In the current study, bats navigating a familiar setting relied on spatial priors to anticipate the perch’s location, directing their sonar—and thus their attention—primarily toward the expected landing region. When the perch was moved, the bats failed to find the perch in the new location, continuing to direct their sonar at the original position. This can explain their failure to land on the perch in its new location, which was just outside their spotlight of attention. These findings suggest that the Egyptian fruit bat’s spotlight of attention is wider when it has access to vision and echolocation together than when it has access to echolocation alone, as animals failed to find the perch-moved by 30 cm in the light and only 15 cm in the dark.

### Multimodal Navigation and the Role of Attentional Focus in Egyptian Fruit Bats

Egyptian fruit bats are well known for their reliance on both vision and echolocation for navigation. Their primary modality appears to be vision, as they exhibit high sensitivity that aids in navigation under scotopic conditions (30,31). However, they have also evolved a lingual echolocation system to complement or substitute for visual information. In the present study, bats showed an increase in sonar click-rates in dark compared to light conditions, consistent with findings from other studies (47,48). Bats also showed a reduction in direct flights toward the perch in darkness, even when the perch was not moved, with direct flights accounting for 42% of the flight trajectories in the light and only 18% in the dark. When visual cues were absent, circling maneuvers allowed bats to refine their flight and landing trajectories, compensating for the loss of visual input by extending time to explore the environment and find the perch. This strategy likely optimizes their approach angle and improves landing accuracy in dark environments (49).

Echolocation, while effective in localizing objects in the dark, demands effective filtering in cluttered environments that give rise to cascades of echo returns from each sonar signal (50–52) and the rapid attenuation of ultrasound limits its operating range (53–57). When both vision and echolocation are available in the light, the perceptual load can be shared across the two sensory systems, with vision providing a longer operating range (33,58). In the light, bats likely used the sensory modality with a longer operating range, i.e. vision, to find the perch. When the perch was displaced by 15 cm, its angular offset from the standard location was small at the bat’s release point of 3 m, and presumably fell within the bat’s spotlight of attention, enabling detection of the perch at its new location. Under light conditions, bats directed their sonar at both the moved perch location and the original location, demonstrating that the bats detected the moved perch, and were able land on it in 86% of trials. When the perch was displaced by 30 cm, the bats successfully landed on the perch in only 39% of the trials, presumably because the angular offset from the standard location fell outside the bat’s spotlight of attention. In the dark, bats could only find the perch using echolocation when they flew closer. This is evidenced by the bats’ failure to find the perch in the dark over hundreds of training trials (see Supplementary Table 1), without first localizing it in the light (see Danilovich and Yovel, 2019, supplementary materials). In the dark, after initial training in the light, the perch was displaced by 15 cm, which is at a larger angular offset from the standard location and fell outside the bat’s spotlight of attention. In the dark, bats landed on the perch displaced by only 15 cm in only 50% of trials.

The bat’s failure to find the moved perch under dark conditions suggests that attentional mechanisms play a crucial role in their navigation. This finding informs a broader debate as to whether inattentional blindness reflects perceptual limitations or memory failures. Research by Ward and Scholl (2015) argues that IB is fundamentally a perceptual deficit, not a memory failure, as their studies demonstrate that individuals can repeatedly fail to perceive salient events in real-time, even when their memory is intact. This is further supported by studies like those of Greene et al. (2017), where high perceptual load tasks led to impaired recognition of unexpected stimuli, suggesting that the attentional system, when overloaded, can lead to IB. Similarly, our study shows that bats operating with sonar in the dark failed to detect the moved perch, indicating a perceptual limitation rather than a memory error. If failed memory were responsible for the bat’s navigation errors, the animals should have performed similarly in the light and dark.

## Conclusions

This study provides evidence for inattentional blindness in a non-human species, Egyptian fruit bats, performing a natural navigation task. Through repeated trials, bats established a spatial prior to find a landing perch. When the perch was unexpectedly moved, established priors interfered with the bat’s ability to find the moved perch, even when visual and echolocation information were available. This behavior aligns closely with the phenomenon of inattentional blindness in humans, where attention to a particular task, cue or location can undermine the detection of salient stimuli. Importantly, this study demonstrates that inattentional blindness can be attributed to perceptual limitations rather than memory failures, as the bats showed large differences in landing success when they had access to two distal sensing modalities, vision and echolocation, as compared with echolocation alone.

This research offers three key advances. First, it reports inattentional blindness in non-human subjects, demonstrating this cognitive phenomenon in an echolocating mammal. Second, it demonstrates that inattentional blindness can occur across different sensory modalities—in this case, vision and echolocation. Third, it offers answers to long-standing questions regarding bat navigation failures, pointing to the dominance of learned spatial priors over sensory perception.

## Methods

### Research Subjects and Experimental Setup

Subjects: Eight Egyptian fruit bats served as subjects in this study and were divided into two groups. The first group consisted of four bats (two males, two females) trained initially in the light (lux level: 28). One female bat died from natural causes before data collection was completed. Of the four bats trained in the light, all met testing success criteria of 80 percent correct landings. Another group consisted of four bats (two males and two females), initially trained in the dark (lux level: 0.76). Information on the training data collected for all bats can be seen in supplementary table 1 (S1 Table). Of the bats trained in the dark, none reached the testing criteria, despite training for multiple days. Bat 08 was trained for only one day because it struggled to navigate in the room. Bat 06 successfully landed on the perch a few times but lacked consistent performance to progress to the testing phase. The bats’ struggle to find the perch in the dark prompted the need to train them initially in the light (S1 Table).

Ethics statement: The handling and treatment of bats adhered to the guidelines outlined by the American Society of Mammalogists (59) and were done in full compliance with the Institutional Animal Care and Use Committee at Johns Hopkins University (protocol number BA23A45). These protocols strictly conformed to the regulations set forth by the Animal Welfare Act and the Public Health Service Policy. The university maintains accreditation by the Association for the Assessment and Accreditation of Laboratory Animal Care International.

Experimental setup: Bats were trained to land on a perch (height: 125 cm) in a set location in the room that was a rest area where food was offered. The flight arena, 5.0m x 2.0m x 2.5m, was lined with acoustic foam on the walls and ceiling, carpeting on the floor, and felt placed over reflective objects to mitigate echoes (Fig 4). In the first phase of training, bats were habituated to the flight room by allowing them to fly freely and naturally find the perch. To control for olfactory cues, food was distributed around the room, but only food on the perch was accessible. Once the bats were accustomed to the flight room and had successfully located the food on the perch, they were released from random locations within the room. A 3-minute interval followed each successful landing to allow the bats the opportunity to rest. The testing phase commenced once the bats reached a success criterion of 80 percent correct landings on the perch for two consecutive days.

**Fig 4:**
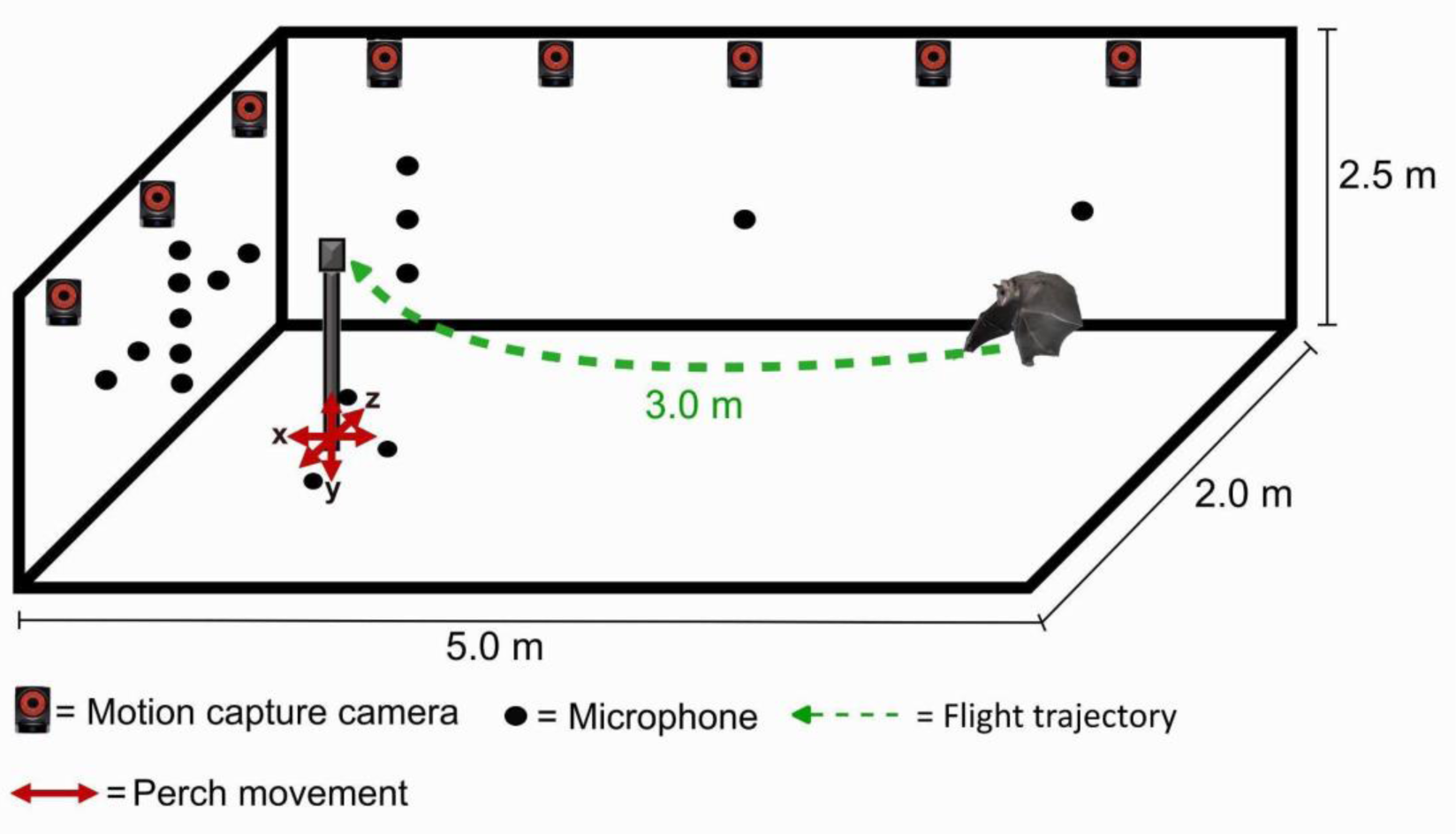
Schematic of test room and navigation task. Schematic of the flight arena showing an example flight trajectory of a bat flying to the perch after being released from the back of the arena. Arrows represent the direction of perch movement (up, down, left, right, front, and back) during testing. A 25-channel microphone array was used to record the directional sonar behavior of the bats (not all microphones are shown). 13 infrared motion capture cameras were used to track the location of the bat as it flew in the room (not all motion cameras are shown).

The training in the light and dark conditions followed the same procedures; however, only bats trained in the light reliably met the success criterion of 80 percent correct landings. The bats trained in the dark were excluded from the experiment, because they failed to reach the 80 percent landing success criterion. Data reported here for bats tested in the dark underwent testing first in the light.

### Experimental Procedure and Design

The experiment was designed to evaluate bat navigation behavior and sonar-guided attention in a standard condition, where the perch was at a fixed location, and in a probe condition, where the location of the perch was moved. During the experiment, bats navigate to a perch location from one of three release locations at the back of the room: Left: One meter from the back wall, 50 cm to the left of the center of the room; Middle: At the back wall, in the center of the room; Right: 50 cm from the back wall, 50 cm to the right of the center of the room. Within an experimental test day, each bat was run in six to eight standard trials, where the perch remained in its original familiar location. Only once per test day was the perch-moved to a different location in the room, and this trial is referred to as a “Perch-Moved” trial. The order of the Perch-Moved trial within the sequence of trials each day was determined following a pseudorandom schedule, with the constraint that the Perch-Moved trial was never the first or last trial of a test day. The trial immediately after the Perch-Moved trial, in which the perch was returned to the standard location, was termed the ‘Post Perch-Moved’ trial. We intentionally limited the number of Perch-Moved trials run per bat, to minimize the bat’s learning about perch location changes. The experiment followed a block schedule design, with each block consisting of four days. On three out of the four days, the perch was moved, averaging three movements per 21 trials. Between blocks, there were one to two days of trials where the perch remained stationary.

### Perch-Moved Conditions

In the Perch-Moved trials, the perch was placed in a new position in one of the following directions—left, right, front, back, up, or down—by either 15 cm or 30 cm (Fig 4). This variation was designed to prevent the bat from predicting the moved perch spatial location. To further prevent the bat from predicting perch movements, the same distance and direction were never repeated within the same block. Occasional equipment failures necessitated the retesting of some conditions and trials. This, along with a deliberately randomized testing schedule and unforeseen events, such as a pregnancy in one of the bats, contributed to variation in the number of trials run with individual bats across conditions. The total number of trials run across conditions for each bat is shown in supplementary table two (S2 Table). The number of Perch-Moved conditions was deliberately limited compared to the standard trials to ensure that the standard position served as the statistical reference point or spatial prior.

### Testing in the Dark

After bats were run in a combined total of 265 trials in the light, an additional 129 trials were run in the dark after bats reached 80% landing success criterion in the dark. This phase of the experiment aimed to determine if bats relied more heavily on spatial priors when using echolocation alone. It also assessed the Egyptian fruit bat’s ability to navigate in the dark after learning the task and the environment in the light. In the dark, the Perch Movement trials were confined to 15 cm, following the same experimental procedures as outlined above. Restricting Perch Movement trials to 15 cm was based on preliminary findings that bats struggled to land on a perch-moved by even this distance in the dark.

### Audio and Video recordings

A 25-channel microphone array of electret condenser microphones (Pettersson Elektronik, Sweden) was used for acoustic data collection (Fig 4). Each microphone was calibrated for sensitivity and directionality using a precision microphone (40DP 1/8”, G.R.A.S., Denmark). Echolocation signals picked up by the microphones were bandpass filtered between 10 kHz and 100 kHz at 10x gain (SBPBP-S1, Alligator Technology, CA) and digitized at sampling frequency of 250 kHz. Three markers placed on microphones in the room were used to record the positions and directions of the microphones in the room. The beam patterns of each sonar click were reconstructed using MATLAB (34), which interpolated the energy spectral density (ESD) of the clicks across all microphones. Corrections for atmospheric absorption and spherical spreading loss were made using recorded temperature and humidity data for each day. Beam aim measurement provided a metric for the animal’s sonar-guided attention.

In each trial, the bat’s 3D flight trajectory was reconstructed using a 13-camera motion capture system (Vicon T40 and T40S, Oxford, UK) operated at a rate of 100 frames per second. Three reflective markers were positioned on a triangle affixed to the bat’s head, which were used to track the bat’s flight trajectory (34). In addition, infrared high-resolution video cameras (Phantom Miro 310, Vision Research, Wayne, NJ, USA), operating at a frame rate of 200, confirmed landing behavior.

## Analysis

### Flight Trajectory

Bat flight trajectories were assigned to one of four outcomes that cataloged the bat’s flight path during each trial: a) direct success: The bat flew directly to the perch and landed on it after release; b) direct failure: The bat approached perch only once but did not successfully land on it; c) back-and-forth success: The bat did not land on the perch on its initial approach but successfully landed after executing more than one approach; d) back-and-forth failure: The bat approached the perch more than once during its flight but failed to land on it.

### Sonar Beam Aim: Metric of Sonar Guided Attention

The directional aim of the bat’s echolocation clicks was used as a metric for the bat’s sonar-guided attention. The directional aim of the bat’s sonar was quantified in relation to the original and moved perch locations. Egyptian fruit bats produce sonar clicks in pairs, typically placing the maximum slope of each click in the pair at the location of a landing perch (60). Clicks within each pair tended to be directed at the same location in the vertical plane in this dataset. In trials where the perch was moved horizontally (i.e., left or right), the direction of the bat’s sonar beam aim was determined by the midpoint of each interclick-pair on the horizontal plane (48). In trials where the perch was moved vertically (i.e., up or down), the beam aim along the vertical plane was determined by the point where the rate of change of energy with height (energy-to-elevation derivative) was nearly at its peak for each click in the pair, which is just above the highest intensity region of the sonar beam (61). The location with the smaller angular distance to the direction of focus was determined to be the point where the bat’s attention was centered, between the standard location and moved perch location (Fig 5). Any sonar signals directed outside of a ±10 deg tolerance of the moved or standard perch locations were categorized as undefined. Details of sonar beam analysis are reported in Lee et al. (2017), which provides a framework for this analysis. Additionally, only trials that contained at least 4 clicks were included for analysis. Some clicks were excluded for analysis because of low SNR or because there was only one click in the click-pair.

### Click-rate

In each trial, the number of clicks emitted by the bat during its initial and subsequent approaches to the perch, as well as the flight times (seconds), travel distances (meters), and the Euclidean distance between the bat and the perch were extracted and plotted in MATLAB version: 9.13.0 (R2022b), (Natick, Massachusetts: The MathWorks Inc.; 2022). The click-rate over time and distance—defined as the number of clicks emitted divided by time (in seconds) and distance (in meters)—for each approach to the perch within a trial was calculated. There was no statistical difference in click-rates calculated over time and distance to the perch so for prosperity purposes only click-rate calculated over distance is presented. Approach 1 was characterized as the period from the bat’s takeoff until it either landed on the perch or passed it during flight. The conclusion of an approach phase was defined by the bat landing on the wall or when the distance from the bat to the perch increased over at least 10 consecutive frames (10 ms). The onset of Approach 2 was determined by a subsequent decrease in the distance to the perch following Approach 1.

Notably, Egyptian fruit bats tend to increase their click-rate in the dark or when preparing to land (35,47). To assess the bat’s click-rate as it approached or closely inspected the perch, we computed the Euclidean distances between the bat and the perch for each emitted click-pair. The bats’ average distance from the perch at which they increased click-rate was then extracted. The average click-rate at this distance was computed across all approaches to the perch, when the perch was in the standard location and when it was moved both 15 cm and 30 cm, in the light and dark.

### Statistical Analysis

All statistical analyses were conducted using R Studio and SPSS. We performed statistical comparisons on independent variables, including bat performance metrics such as *Flight Trajectories* and *Landing Success*, as well as indices of sonar-guided attention, specifically the *Direction of Sonar Beam Aim* and *Click-Rate*. These independent variables were modeled against dependent variables such as *Perch Displacements* (Standard, Moved, Post-Moved), *Perch Displacement Directions* (up, down, left, right, back, front), *Lighting* (light, dark), *Release Position* (left, right, center), *Individual Bat Subjects*, and *Time Periods* (across days).

### Flight Trajectories

A Multinomial GLMM with Repeated Measures was used to evaluate the effects of lighting condition, perch displacement, and individual bat subjects on flight trajectories, which had four possible outcomes: direct success, back-and-forth success, direct failure, and back-and-forth failure. Post-hoc pairwise comparisons with Estimated Marginal Means (EMM), were done to assess interactions between independent variables. Bonferroni corrections were applied to control for multiple comparisons and to minimize Type I error.

### Landing Success

Landing Success and Flight Trajectories were modeled against all listed independent variables to examine how factors such as perch displacement, lighting conditions, and release positions influenced bats’ ability to find and navigate to a perch. Generalized Linear Mixed Models (GLMMs) with repeated measures were used with a binary logistic regression to account for repeated measures, as each bat was subjected to multiple trials across different conditions. GLMMs were followed by post-hoc Tukey Unequal-N tests with Bonferroni correction to control for multiple comparisons and reduce the likelihood of Type I errors.

### Sonar Beam Aim

A Kruskal Wallis test was used to investigate where bats directed their sonar attention when the perch was moved in both light and dark. The average number of click-pairs (dependent variable) directed at the standard, moved, or at undefined locations was compared across different independent variables including perch displacement (15 cm and 30 cm) and lighting condition (light or dark). Dunn’s post-hoc test with Benjamini-Hochberg adjustment was applied to identify specific pairwise differences when significant results were observed.

### Click-rate (calculated over distance, and near the perch)

Generalized Linear Mixed Models (GLMMs) were used to model the relationship between click-rates—over the approach to the perch and near the perch—across lighting conditions, perch displacements and landing success. Separate GLMMs were fitted for each click-rate variable, with lighting condition, perch displacement, and landing success and their interactions included as fixed effects, and individual bat as a random effect to account for individual variability. Post-hoc pairwise comparisons were performed using estimated marginal means (EMM) to identify specific differences between levels of lighting conditions, perch displacement, and landing success. Pairwise comparisons were made between bats that landed and did not land across different lighting and perch displacement conditions. Type III ANOVAs were used to evaluate the significance of each factor and interaction in the models. Bonferroni corrections were applied to control for multiple comparisons and reduce the likelihood of Type I errors. Additionally, a separate Friedman ANOVAs followed by a Wilcoxon Signed Rank Test was used to test the difference in click-rates within each bat navigating in the dark and light in control trials (no perch movement), where the bats reached 80 % success criteria, across all approaches to the perch.

## Acknowledgments

We thank Dimitri Skandalis who gave valuable feedback in the early stages of experimental conceptualization. We thank Jim Garmon for designing and manufacturing the headpiece that was used to track the bats’ 3D positioning and head aim. We thank Davi Drieskins and Clarice Diebold for their help with the figures. Thank you to the undergraduate researchers who contributed to data collection Alara Kaplanoglu, Leslie Bucio, and Marianna Meade. We also thank the students who contributed to data analysis Alara Kaplanoglu, Leslie Bucio, Marianna Meade, Laura Panlilio, Myles Gosha, Ruya Ozveren, and Sadie Friesen.

## Funding Acknowledgement

We gratefully acknowledge fellowship support from the Center for Hearing and Balance T32 Postdoctoral Fellowship T32DC000023-39 (N.M.F.). This research was supported by the following research grants: NIH R01 NS121413 (C.F.M.), Human Frontiers Science Program Research Grant RGP0045/2022 (C.F.M.), NSF Brain Initiative Grant NCS-FO 1734744 (C.F.M.), NSF CRCNS Grant 2011619 (C.F.M.), and Office of Naval Research Grants N00014-23-1-2086 and N00014-17-1-2736 to C.F.M.

## Notes

### Competing Interest Statement

The authors have declared no competing interest.

