## Supplementary for "Why do bats fly into cave doors? Inattentional blindness in echolocating animals"

**Supplementary material**

| | | Light (# of trials) | | | | Dark (# of trials) | | | |
|---|---|---|---|---|---|---|---|---|---|
| Initial Training | **Bat** | **1*** | **2*** | **3*** | **4*** | **5** | **6** | **7** | **8** |
| | **Total Training** | 72 | 9 | 43 | 39 | 39 | 295 | 57 | 12 |
| | **First landing on perch** | 17 | 2 | 15 | 13 | | 13 | | |
| Retraining in the dark | **Bat** | **1*** | **2*** | **3*** | | | | | |
| | **Total Training** | 26 | 0 | 32 | | | | | |
| | **First landing on perch** | 17 | 0 | 27 | | | | | |

**S1 Table: Summary of Training Data:** Left columns show the number of training trials required before individual bats reached criterion performance of 80% successful landing in the light and retraining in the dark. Asterisks show which bats met this criterion. First landing on the perch is the number of trials recorded before each bat successfully landed on the perch for the first time. On average, bats received 6-8 training trials per day, mirroring the number of trials bats received during testing sessions. Right columns show the number of trials run initially in the dark. No bat reached the 80% successful landing criterion under this condition, and only one bat landed on the perch at all.

| | | Standard | | | | Perch-moved 15 cm | | | | Perch-moved 30 cm | | | | Post-moved | | | |
|---|---|---|---|---|---|---|---|---|---|---|---|---|---|---|---|---|---|
| | Bat | Land | Fail | Total | SR (%) | Land | Fail | Total | SR (%) | Land | Fail | Total | SR (%) | Land | Fail | Total | SR (%) |
| Light | 1 | 36 | 1 | 37 | 97 | 6 | 0 | 6 | 100 | 1 | 3 | 4 | 25 | 9 | 1 | 10 | 90 |
| | 2 | 77 | 2 | 79 | 97 | 8 | 2 | 10 | 80 | 3 | 4 | 7 | 43 | 19 | 0 | 19 | 100 |
| | 3 | 60 | 3 | 63 | 95 | 6 | 1 | 7 | 86 | 3 | 4 | 7 | 43 | 11 | 3 | 14 | 79 |
| Dark | 1 | 28 | 9 | 37 | 76 | 2 | 3 | 5 | 40 | | | | | 0 | 3 | 3 | 0 |
| | 2 | 33 | 2 | 35 | 94 | 3 | 1 | 4 | 75 | | | | | 4 | 0 | 4 | 100 |
| | 3 | 26 | 11 | 37 | 70 | 2 | 1 | 3 | 67 | | | | | 2 | 1 | 3 | 67 |

**S2 Table: Landing Performance Across Experimental Conditions.** Summary of the landing success and failure for each bat under three experimental conditions: standard, perch-moved 15 cm, and perch-moved 30 cm and the trial after the perch was moved. For each condition, the total number of trials and the percentage success (SR%) are displayed. The data is divided into two conditions: initial testing in light and retesting in dark. The perch-moved 30 cm condition in the dark was not tested.

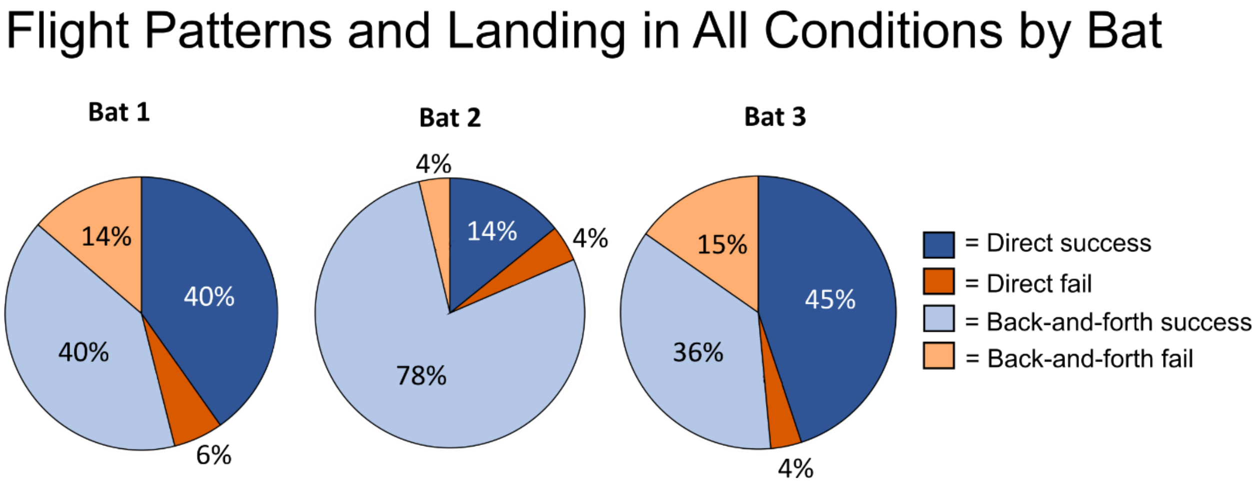


**S3 Figure: Flight Patterns and Landing of Individual Bats Across All Conditions.** Pie charts show the flight patterns of each bat across all testing conditions. Bat 2 predominantly exhibited back-and-forth flights, whereas Bats 1 and 3 displayed a more varied distribution of flight patterns.

| | Bat | Standard (avg. click pairs) | | | Perch-moved 15 cm (avg. click pairs) | | | Perch-moved 30 cm (avg. click pairs) | | |
|---|---|---|---|---|---|---|---|---|---|---|
| | | Familiar | Moved | Undefined | Familiar | Moved | Undefined | Familiar | Moved | Undefined |
| Light | 1 | 3.6 | | 0.8 | 2 | 2.5 | 1.5 | 3 | 0.3 | 2.3 |
| | 2 | 2.8 | | 1.7 | 5 | 0 | 1 | 3.5 | 1 | 1.5 |
| | 3 | 4.8 | | 1.9 | 1 | 2 | 2 | 5 | 0 | 1 |
| Dark | 1 | 4.5 | | 3.0 | 5 | 0 | 2.3 | | | |
| | 2 | 2.4 | | 1.9 | 1 | 2 | 2 | | | |
| | 3 | 6.4 | | 2 | 2 | 1 | 3 | | | |

**S4 Table: Beam Aim Direction to Perch in Light and Dark Tests.** Average number of click pairs directed at the standard location, and new location after the perch was displaced by 15 cm and 30 cm from the standard location. No data were collected for the perch-moved 30 cm condition in the dark. A click pair directed outside the two locations of interest were categorized as undefined, based on a ±10 deg tolerance to the nearest perch location. This table shows that bats tested in the light tended to direct their sonar beam toward the standard and moved locations after the perch was displaced by 15 cm but predominantly toward the standard location after the perch was displaced by 30 cm. Bats tested in the dark tended to direct their sonar towards the standard location after the perch was displaced by 15 cm.

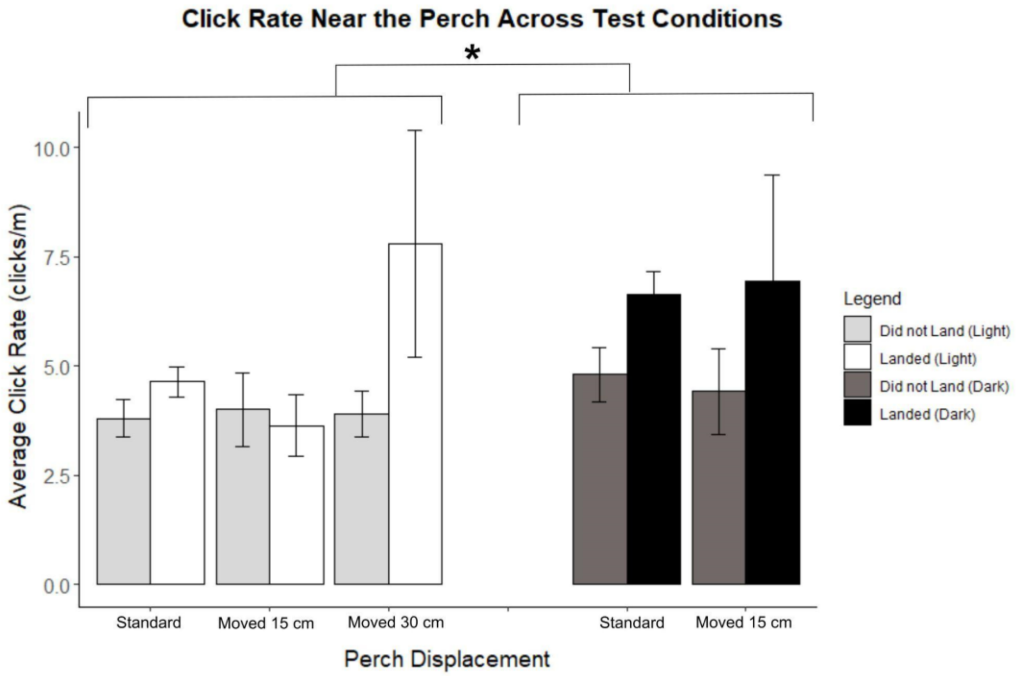


**S5 Figure: Echolocation Click Rates Near the Perch.** Bar graphs depicting the average echolocation click-rate (±SE) of bats within a 1.5-meter radius of the perch when it did and did not land, compared across perch positions in both light and dark conditions. Sonar click-rate did not significantly differ between landing and not landing, though in the 30 cm perch displacement condition and in the dark bats showed higher click rates when they landed.

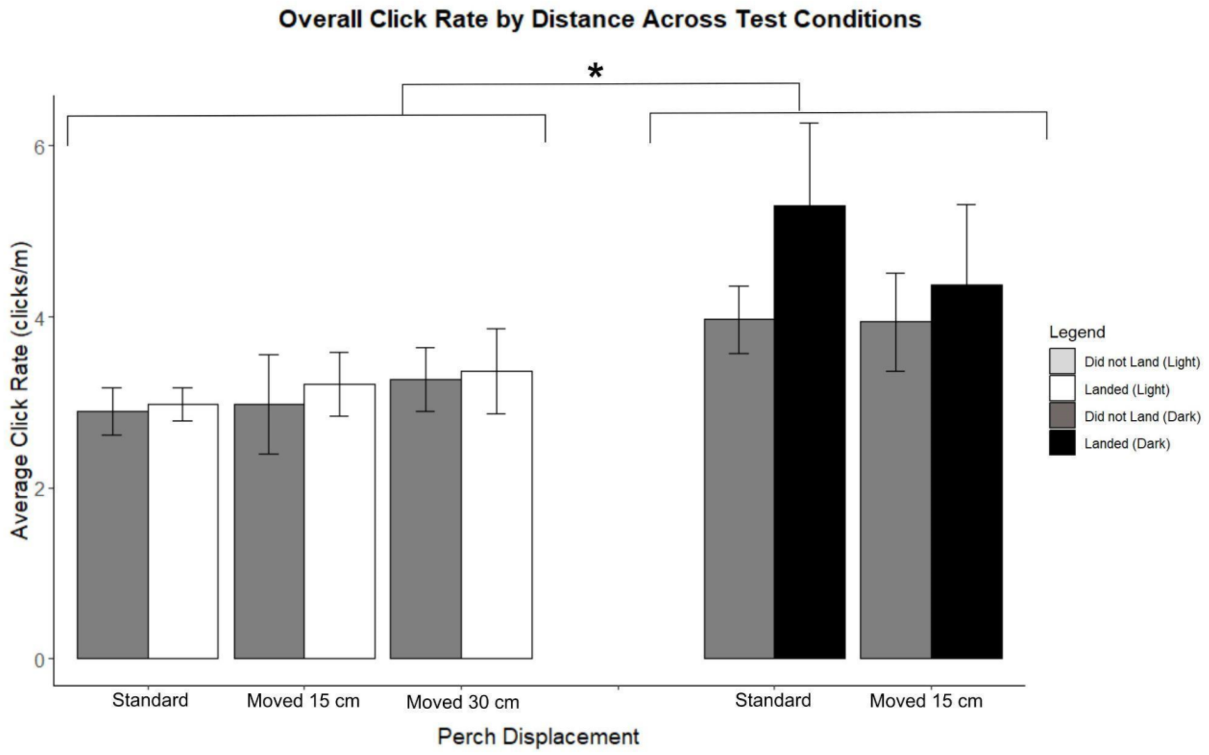


**S6 Figure: Echolocation Click Rates Over All Testing Conditions.** The average click rate (± SE) of bats is shown, separated by light versus dark conditions and whether the bats landed or did not land. Bats produced significantly more sonar clicks in the dark compared to the light. Asterisk indicates significance.
